# Structural characterization of an intrinsically disordered protein complex using integrated small-angle neutron scattering and computing

**DOI:** 10.1101/2022.12.19.521134

**Authors:** Serena H. Chen, Kevin L. Weiss, Christopher Stanley, Debsindhu Bhowmik

## Abstract

Characterizing the structural ensembles of intrinsically disordered proteins (IDPs) is essential for studying structure-function relationships as conformational dynamics govern proteins’ biological functions. Due to the notable difference between the neutron scattering lengths of hydrogen and deuterium, selective labeling and contrast matching in small-angle neutron scattering (SANS) becomes an effective tool to study dynamic structures of disordered systems. However, the experimental timescale typically results in measurements averaged over multiple conformations, leaving complex SANS data for disentanglement. We hereby demonstrate an integrated method to elucidate the structural ensemble of a protein complex formed by two IDP domains, the NCBD/ACTR complex, using data from selective labeling SANS experiments, microsecond all-atom molecular dynamics (MD) simulations with four molecular mechanics force fields, and an autoencoder-based deep learning (DL) algorithm. By incorporating structural metrics derived from the SANS experiments as constraints in MD structure classification, we describe a structural ensemble that captures the experimental SANS and, in addition, NMR data better than MD ensembles generated by one single force field. Based on structural similarity, DL reveals three clusters of distinct conformations in the ensemble. Our results demonstrate a new integrated approach for characterizing structural ensembles of IDPs.

## Introduction

Prevalent across the three domains of life, intrinsically disordered proteins (IDPs) are important for a wide range of biological functions from molecular recognition to regulation of enzyme activity^1-4^. As the function of a protein depends on its structural dynamics, it is inevitable to study the structural ensembles of IDPs to uncover their physiological roles. Yet, due to their extensive structural flexibility, determining structural ensembles of IDPs presents a challenge for both experiment and computation.

A variety of experimental techniques have been developed and employed to characterize structural ensembles of IDPs, including innovative variations of nuclear magnetic resonance (NMR) spectroscopy^5, 6^ and single-molecule approaches^7, 8^. Complementary to these techniques, small-angle X-ray and neutron scattering (SAXS and SANS) offer global structural information of IDPs in terms of shape and size which are useful for validation of high-resolution structures^9-11^ and elucidating complexes^12^. Compared with X-rays, neutrons are scattered by atomic nuclei, leading to their high sensitivity to light elements, such as hydrogen. Different hydrogen isotopes, such as hydrogen and deuterium, result in distinct neutron scattering lengths. Therefore, by selective deuteration and varying the D_2_O to H_2_O ratio of the solvent, one can modulate contrast between different components of a system, making SANS an ideal probe for studying structural ensembles of IDPs^13-17^. While SANS contrast matching and variation experiments typically use selective deuteration of an entire component, methods of selective labeling at residue level were developed to characterize dynamics within globular proteins^18-20^. Here we employed these capabilities to measure an IDP complex involving a residue-specific deuterium-labeled protein, with the remaining hydrogenated portion contrast matched out. To the best of our knowledge, these methods have not been demonstrated in IDPs before.

Due to the experimental timescale, SANS, similar to other ensemble methods, yields scattering intensities averaged over all conformations. In this respect, computational techniques complement experiment by modeling a pool of conformations to describe experimental data. Modeling is predominately based on three approaches. One approach is to apply the experimental data as restraints for conformational sampling^21, 22^. Structural ensembles constructed by this approach rely on the number of experimental restraints as well as how stringent the restraints are enforced when sampling. Another approach, used by the ensemble optimization method (EOM)^23^ and ENSEMBLE^24^, is to generate a large number of possible conformations from the conformational space, and then select a subset of the conformations which fit the experimental data. The outcomes of this approach depend on the quality and diversity of the initial pool of conformations. The other approach applies various enhanced sampling techniques to cross free energy barriers and sample more conformational space^25, 26^. This approach remains computationally intensive and requires careful scrutiny for convergence^27^.

Here we integrated residue-specific deuterium labeling of SANS, molecular dynamics (MD) simulation, and an autoencoder-based deep learning (DL) algorithm to study the structural ensemble of a protein complex formed by two IDP domains, the nuclear co-activator binding domain (NCBD) of CREB binding protein and the activation domain of the p160 transcriptional co-activator for thyroid hormone and retinoid receptors (ACTR). The NCBD/ACTR complex regulates gene expression and has a long-known association with breast and ovarian cancers^28^. The complex formation, determined by NMR spectroscopy, illustrated the first example of coupled binding and folding of IDPs^29^. While the NCBD/ACTR complex is more rigid than their unbounded states, 20% of the residues of the complex remain highly flexible^30^. To gain insights into dynamic structures of the NCBD/ACTR complex, we performed microsecond all-atom MD simulations with four molecular mechanics force fields and compared the structural ensembles generated by these force fields. To study the effect of different experimental constraints on the structural ensemble, we classified MD structures into ensembles based on structural metrics derived from SANS experiments that used both full contrast and residue-specific labeling with contrast matching. We then verified the ensembles by recapitulating scattering intensities and NMR chemical shifts. Lastly, we characterized the composition of distinct conformations populated in a representative ensemble that showed the best agreement with the experiments by a convolutional variational autoencoder. Our work provides an integrated SANS and MD/DL workflow for characterizing structural ensembles of IDPs.

## Results and Discussion

To conduct selective labeling SANS experiments, we deuterated the mouse NCBD peptide at all five alanine (Ala) and seven leucine (Leu) positions (*d*_A,L_-NCBD) (**Figure 1A**) and collected SANS data using the extended Q-range small-angle neutron scattering instrument at the Spallation Neutron Source located at Oak Ridge National Laboratory^31^. Ala and Leu residues were chosen based on commercially available FMOC deuterated amino acids for the peptide synthesis. The motivation to label all of these positions was to provide sufficient signal for the SANS experiment. By adjusting the buffer solution to 40% D_2_O for *d*_A,L_-NCBD/ACTR, the hydrogenated portion is suppressed such that the measured signal predominately reflects NCBD *d*-Ala and *d*-Leu correlations. As a control measurement, SANS was performed on hydrogenated NCBD (*h*-NCBD/ACTR) in 40% D_2_O buffer, where a linear-linear plot of the scattering profile illustrates the contrast matched complex compared to the signal from *d*_A,L_*-*NCBD/ACTR in the same 40% D_2_O buffer (**Figure 1B)**. Indeed, the calculated match point of the *d*_A,L_-NCBD/ACTR hydrogenated component (i.e., excluding *d*-Ala and *d*-Leu residues) is 43.7% D_2_O using MULCh_32_. A full contrast SANS profile also was obtained for all-hydrogenated, *h*-NCBD/ACTR in 100% D_2_O buffer. After initial Guinier fits to the SANS data (**Figures S1A, S1B**), pair-distribution, *P*(*r*), profiles were calculated yielding the radius of gyration (*R*_g_) for each case (**Figure 1C**). The linearized Guinier plots and fits provide a quality assessment of the low Q portion of the scattering curves, while the *P*(*r*) analysis covers a broader Q range of the data. Based on the differences between the Guinier and *P*(*r*) fit methods, slight discrepancies in the obtained *R*_*g*_ values can be expected, as we note particularly for the *d*_A,L_-NCBD/ACTR in 40% D_2_O case (see **Table S1**). We also investigated the magnitude of the dependence of *R*_*g*_ values on chosen *D*_*max*_ in the *P*(*r*) fitting procedure. We performed multiple *P*(*r*) fits with varying *D*_*max*_, while staying within an appropriate range to still obtain good fits (see Figure 1C). The *R*_*g*_ values are consistent; they only vary by ∼0.5 Å in both cases and by no more than 0.8 Å if the extremes between error bars are considered (**Figures S1C, S1D**). For all of our further analyses, we used representative *P*(*r*) fit results, where the *R*_*g*_ values of *h*-NCBD/ACTR in 100% D_2_O and *d*_A,L_-NCBD/ACTR in 40% D_2_O are 15.0 ± 0.1 Å and 11.7 ± 0.1 Å, respectively. These *R*_*g*_ values were used as constraints to classify MD structures of the NCBD/ACTR complex into ensembles for further characterization.

**Figure 1.**
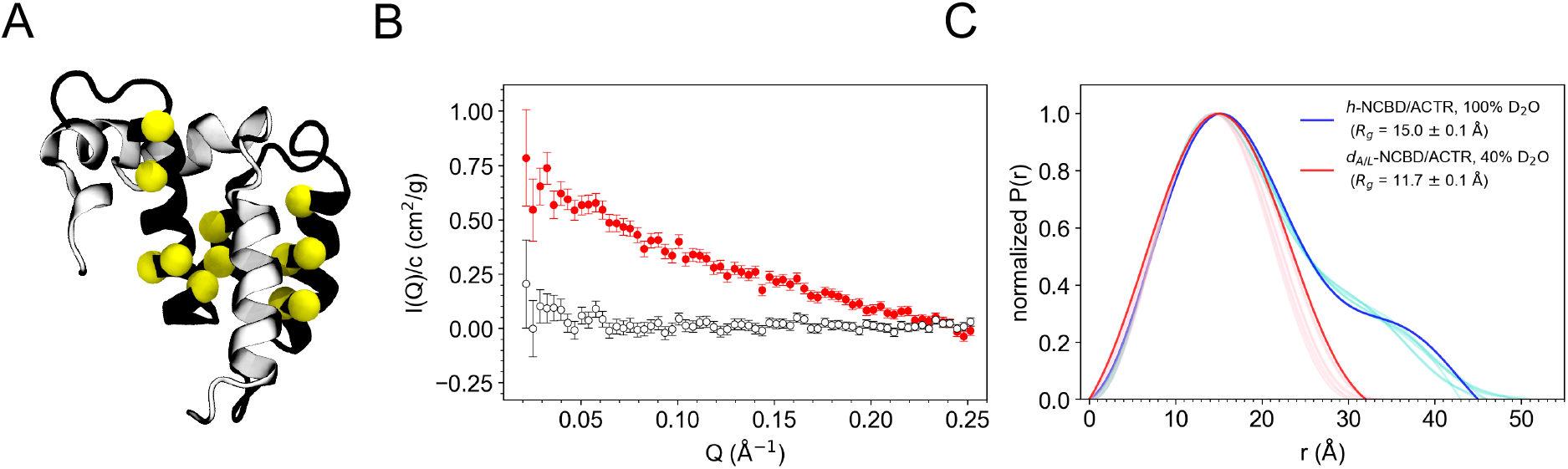
Selective deuteration and SANS provide representative *R*_*g*_ values for NCBD/ACTR structural characterization. (A) Model of the NCBD/ACTR complex illustrating the selectively deuterated Ala and Leu residues of NCBD used for neutron experiments. NCBD (black ribbon) was selectively deuterated at all five Ala and seven Leu positions (yellow spheres). ACTR (gray ribbon) is also shown. (B) Linear-linear plot of SANS curves to illustrate the contrast matched NCBD/ACTR control experiment. SANS curves are normalized by concentration, *I*(*Q*)/*c*. The all-hydrogenated, *h*-NCBD/ACTR complex at 40% D_2_O (black, open symbols) is contrast matched, showing no scattering signal above background. The measurable scattering from selectively labeled, *d*_A,L_-NCBD/ACTR in 40% D_2_O (red, solid symbols) is shown for comparison. (C) Pair-distribution profiles calculated from the SANS data for *h*-NCBD/ACTR in 100% D_2_O (blue) and *d*_A,L_-NCBD/ACTR in 40% D_2_O buffer (red). Additional profile calculations, with varying *D*_*max*_, are also shown for each in light blue and red, respectively (also see Figures S1C, D). The *R*_*g*_ values from the representative profiles are listed in the legend (also see Table S1).

To compare with experimental data, we generated a comprehensive pool of structures using multiple atomistic molecular mechanics force fields in MD simulations and explored the effect of force fields on simulated NCBD/ACTR complex structures^33, 34^. We started from an NMR model of PDB 1KBH^29^ and performed a total of 12 µs all-atom MD simulations in explicit solvent with four widely used Amber and CHARMM force fields: a99SB^35^, a99SB-ILDN^36^, a99SB-*disp*^37^, and C36m^38^. The a99SB and a99SB-ILDB force fields were developed primarily to target folded proteins. The a99SB-*disp* and C36m force fields are more recent updates, which address both folded and disordered proteins. Although some studies show deuteration effects in force field parameters^39^ as well as on protein stability and dynamics^40^, the computationally rigorous approach of reparametrizing *d*-Ala and *d*-Leu with D_2_O solvent is beyond the scope of this work. Instead, here, our focus is to study structural ensembles of unmodified proteins in the native environment, i.e., H_2_O solvent, and understand how this compares to contrast matching SANS experimental data. Toward this aim, we performed MD simulations using unmodified hydrogen mass and force field parameters with H_2_O solvent. To assess structural flexibility at the deuterated positions, Ala and Leu of NCBD, in our simulations, we highlighted the positions of the deuterium labeled side chains in a representative MD trajectory and calculated the average C_*α*_ root mean-square fluctuation (RMSF) per residue of ACTR and NCBD structures in four MD ensembles (**Figure S2**). In all MD structures, the deuterated positions show little fluctuation (less than 2 Å), suggesting that these sites are relatively stable. Moreover, we computed helical fraction and all-atom root mean-square deviation (RMSD) of overall MD structures from the NMR model (**Figure S3**). The MD structures sampled by different force fields have distinct helical fraction distributions in ACTR and NCBD, especially in the first 30 residues of ACTR. Overall, the a99SB structures have the lowest average helical content and the C36m structures have the highest average helical content. Interestingly, the regions of low helical fraction also correspond to regions of higher RMSF values in Figure S2. However, the average helical fraction of these regions is not zero, suggesting flexible movement of ordered helices and helix-loop conversion in the complex. RMSD distributions of the MD structures range from 4 to 10 Å, suggesting various degrees of flexibility within the complex structure. The trends of the median RMSD and average helical fraction of the four forcefields are reversed, where the C36m structures have the lowest median RMSD value and the highest helical fraction, and the a99SB structures have the highest median RMSD value and the lowest helical fraction. Interestingly, despite their higher helical content, the a99SB-*disp* structures have the broadest RMSD distribution, followed by the C36m structures. The RMSD distributions of the a99SB and a99SB-ILDN structures are relatively narrow.

Next, to evaluate the agreement between MD simulations and experiments, we compared MD-derived and experimental SANS curves and NMR chemical shifts. For each MD NCBD/ACTR complex structure we computed two SANS scattering curves and associated *R*_*g*_ values following the same deuteration pattern in the experiments using the CRYSON program^41^ without experimental fitting to prevent bias. The scattering curves overlap well with the experiments, especially at low Q, suggesting that all MD ensembles have similar average shape (**Figure 2A**). To further compare theoretical and experimental scattering curves at different Q range, we calculated the reduced chi-square value (χ^2^) against increasing Q range using eq. (1).

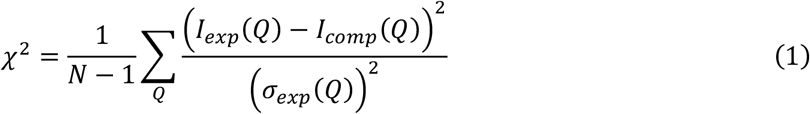

*I*_*exp*_ and *I*_*comp*_ are the scattering intensity measured by SANS experiments and by computation, respectively. *σ*_*exp*_ is the standard deviation of the experiments. *N* is the total number of structures in an ensemble. There are 27,000 structures in each MD ensemble. The lower bound of all Q ranges is 0.02 Å^−1^, and the upper bound increases from 0.03 Å^−1^ to 0.30 Å^−1^ for *h*-NCBD/ACTR in 100% D_2_O and from 0.03 Å^−1^ to 0.25 Å^−1^ for *d*_A,L_-NCBD/ACTR in 40% D_2_O, both with an interval of 0.01 Å^−1^. The result of the χ^2^ value computed across the increasing Q range is summarized in **Figures 2B, 2C**. Even though all MD ensembles fit well at the low Q, for *h*-NCBD/ACTR in 100% D_2_O, the a99SB*-disp* ensemble has the lowest χ^2^ value especially when Q is less than 0.15 Å^−1^. At the high Q, the χ^2^ value increases due to the background scattering, but overall, the C36m ensembles has the lowest χ^2^ value when Q is greater than 0.15 Å^−1^ . The χ_2_ value for *d*_A,L_-NCBD/ACTR in 40% D_2_O is small for all ensembles, which ranges between 1.18 and 2.18 for the largest Q range, from 0.02 Å^−1^ to 0.25 Å^−1^. The small χ^2^ values support the observation found in the MD simulations that the deuterated positions are relatively stable.

**Figure 2.**
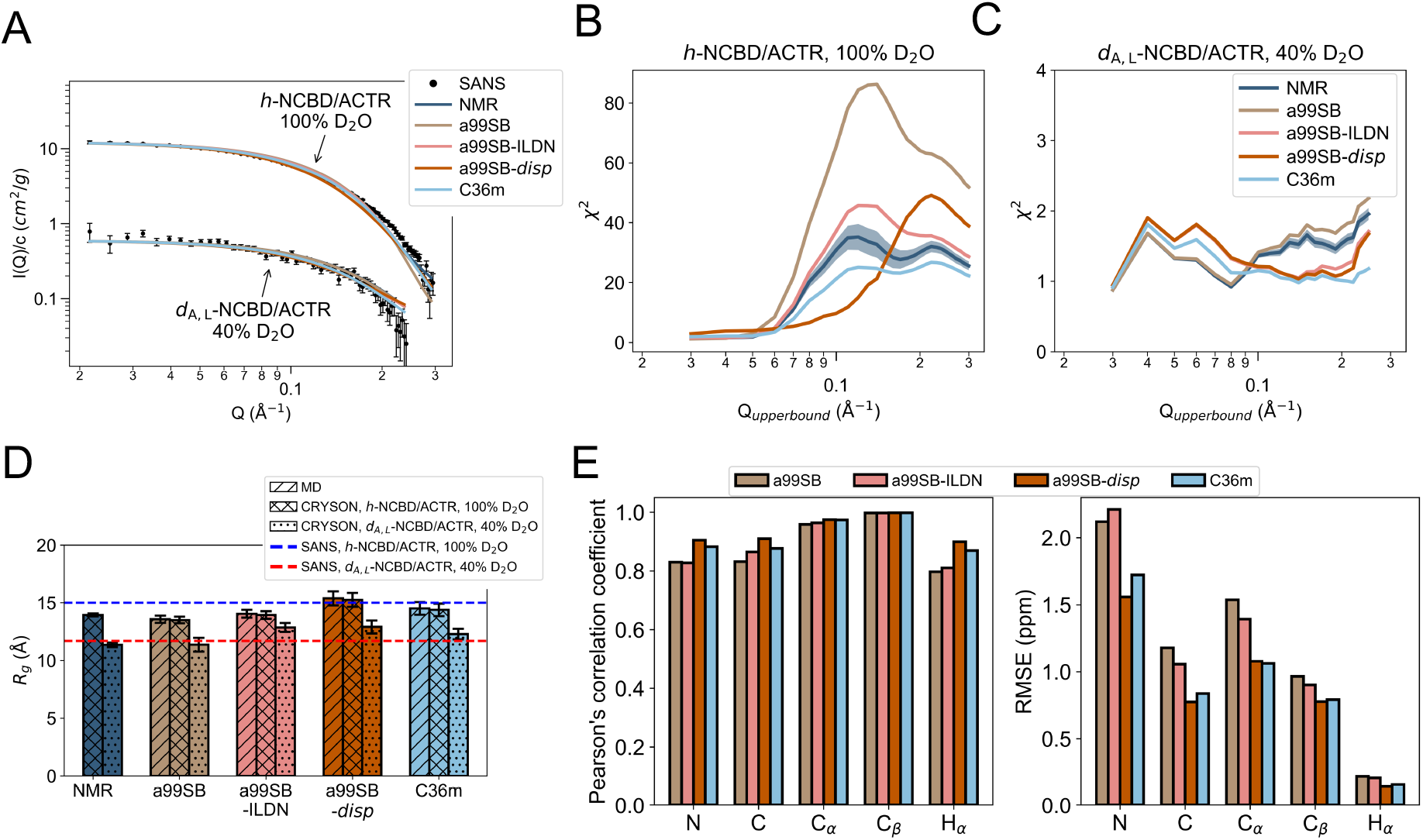
Comparison of NCBD/ACTR force field-based structural ensembles from MD simulations reveals that a99SB-*disp* and C36m show the strongest agreement with SANS and NMR experiments. (A-C) Comparison of calculated average SANS curves by CRYSON^41^ to experimental SANS curves. (A) SANS curves normalized by concentration, I(Q)/*c*, against Q from 0.02 Å^−1^ to 0.30 Å^−1^ for *h*-NCBD/ACTR in 100% D_2_O and against Q from 0.02 Å^−1^ to 0.25 Å^−1^ for *d*_A,L_-NCBD/ACTR in 40% D_2_O. The experimental SANS data are shown as black points, where error bars represent the standard deviation. The computed average SANS curves are depicted as lines, where NMR is in navy, a99SB in tan, a99SB-ILDN in pink, a99SB*-disp* in dark orange, and C36m in light blue. Note that the standard deviations of the computed SANS curves are shown but lie within the line plots. (B, C) Reduced chi-square values (χ^2^) of the SANS curves between experiments and each of the NMR and MD ensembles for (B) *h*-NCBD/ACTR in 100% D_2_O and (C) *d*_A,L_-NCBD/ACTR in 40% D_2_O. The χ^2^ is plotted against increasing Q range, in which the lower bound is 0.02 Å^−1^ and the upper bound (Q_upperbound_) is from 0.03 Å^−1^ to 0.30 Å^−1^ in B and to 0.25 Å^−1^ in C, with an interval of 0.01 Å^−1^. Shaded error bars represent 95% confidence interval of the mean. Note that the error bars of the χ^2^ for the MD ensembles are shown but most of them lie within the line plots. (D) Radius of gyration (*R*_*g*_) of MD structures are computed using atomic coordinates and masses in GROMACS_42_ (MD) as well as using CRYSON with and without selective deuteration (CRYSON *d*_A,L_-NCBD/ACTR in 40% D_2_O vs. CRYSON, *h*-NCBD/ACTR in 100% D_2_O). The average *R*_*g*_ values of the twenty NMR structures using CRYSON with and without selective deuteration are included for comparison. The *R*_*g*_ values derived from the experimental SANS curves of *h*-NCBD/ACTR in 100% D_2_O and *d*_A,L_-NCBD/ACTR in 40% D_2_O are shown as dashed lines in blue and red, respectively. Error bars represent the standard deviation among the structures of each ensemble. (E) Comparison of MD-derived and experimental chemical shifts by (left) Pearson’s correlation coefficients and (right) RMSEs. Chemical shifts of MD structures are computed using SPARTA+^43^. Unless mentioned otherwise, all analyses are performed on twenty NMR structures and 27,000 MD structures per ensemble (three independent trajectories of 9,000 structures each, with the first 100 ns/1,000 frames discarded).

We also compared the *R*_*g*_ values of MD structures computed using GROMACS_42_ as well as using CRYSON with and without selective deuteration (**Figure 2D**). As expected, the *R*_*g*_ values calculated by GROMACS and CRYSON without selective deuteration are comparable. The a99SB*-disp* and C36m structures have the *R*_*g*_ values that are the closest to the SANS experiments in both sample conditions. To further characterize individual NCBD/ACTR complex structures of each MD ensemble, we grouped the MD structures into four categories using their two calculated *R*_*g*_ values: “100_+_40_+_”, “100^+^40^-^”, “100^-^40^+^”, and “100^-^40^-^” (**Figure S4**). A structure fell into the “100^+^40^+^” category if both calculated *R*_*g*_ values are within the two experimental *R*_*g*_ constraints listed in Figure 1C. If only one of the *R*_*g*_ values satisfies the experimental constraints, the structure is in the “100_+_40_-_” or “100_-_40_+_” category. If neither calculated *R*_*g*_ values satisfies the experimental constraints, the structure is in the “100_-_40_-_” category. As a comparison, we also computed the *R*_*g*_ values of all twenty NMR models in PDB 1KBH. Surprisingly, none of the twenty NMR structures satisfies both experimental constraints. Three out of the twenty structures belong to 100^-^40^+^, and the other seventeen structures are 100^-^40^-^. A similar distribution is observed in the a99SB structural ensemble. About 12.7% of the NCBD/ACTR structures are 100^-^40^+^, and the rest are mostly 100^-^40^-^. (There is one 100^+^40^-^structure in one of the three replicas near 950 ns.) As the NMR structures are refined using an Amber force field^29^, it is likely that these early versions of Amber force fields are biased to the 100^-^40^+^ and 100^-^40^-^conformations. The a99SB-ILDN force field improves the accuracy of the side chain torsion potentials for four amino acids of the a99SB force field^36^. However, these modifications do not improve sampling of NCBD/ACTR structures that satisfy the constraints. Less than 0.1% of the sampled structures are 100^+^40^-^or 100^-^40^+^, while most of the sampled structures satisfy neither experimental constraint. Compared with the a99SB and a99SB-ILDN structural ensembles, the a99SB-*disp* force field generates the NCBD/ACTR complex structures in all four categories. About 0.2% of the NCBD/ACTR complex are 100^+^40^+^, 17.7% are 100^+^40^-^, 1.1% are 100^-^40^+^, and the remaining 81.0% are 100^-^40^-^. Similarly, the C36m force field also generates a heterogeneous ensemble of the NCBD/ACTR complex in all four categories. The C36m distribution of 100^+^40^+^, 100^+^40^-^, 100^-^40^+^, and 100^-^40^-^structures are 0.7%, 7.3%, 5.6%, and 86.4%, respectively. Although both the a99SB-*disp* and C36m force fields generate a small amount of the 100^+^40^+^ structures, they appear mostly after 800 ns in the a99SB-*disp* ensemble while they are distributed consistently throughout 1 µs in the C36m ensemble. According to this initial comparison of the MD-derived SANS curves and *R*_*g*_ values with the experimental SANS data, we found that the C36m and a99SB-*disp* force fields describe the NCBD/ACTR complex structure better than the a99SB and a99SB-ILDN force fields. The same trend applies when we compared MD-derived and experimental chemical shifts of different nuclei, where the C36m and a99SB-*disp* structures show the strongest agreement with the experiments with overall the highest correlation coefficients and the lowest RMSE values (**Figure 2E**).

To evaluate the effect of *R*_*g*_–based structural classification on the ensemble, we compared the 100^+^40^+^, 100^+^40^-^, 100^-^40^+^, and 100^-^40^-^ensembles to the SANS and NMR experiments. There are 233, 6772, 5265, and 95750 structures in the 100^+^40^+^ 100^+^40^-^, 100^-^40^+^, and 100^-^40^-^ensembles, respectively. **Figures 3A-3C** illustrate their computed average SANS curves and χ^2^ values against increasing Q range as compared to experimental SANS curves. The 100^+^40^+^ ensemble has the lowest χ_2_ value at all Q ranges compared with the other *R*_*g*_–based ensembles, suggesting that the 100_+_40_+_ ensemble captures shape and size that best describes experimental SANS curves. As a validation, we calculated average *R*_*g*_ value of each *R*_*g*_–based ensemble and confirmed the procedure of grouping each of the NMR and MD structures into one of the four categories based on the two calculated *R*_*g*_ values described above (**Figure 3D**). In addition, we repeated the analysis of comparing computed chemical shifts of each *R*_*g*_–based ensemble with experimental chemical shifts (**Figure 3E**). The 100_-_40^-^and 100^-^40^+^ ensembles have slightly lower correlation coefficients and higher RMSE values for all nuclei, which is expected as the two ensembles mainly consist of a99SB and a99SB-ILDN structures (**Figure S4B**). However, comparing Figures 2E and 3E, we found that the *R*_*g*_–based ensembles show better agreement with experiments than the force field-based ensembles with overall higher correlation coefficients and lower RMSE values.

**Figure 3.**
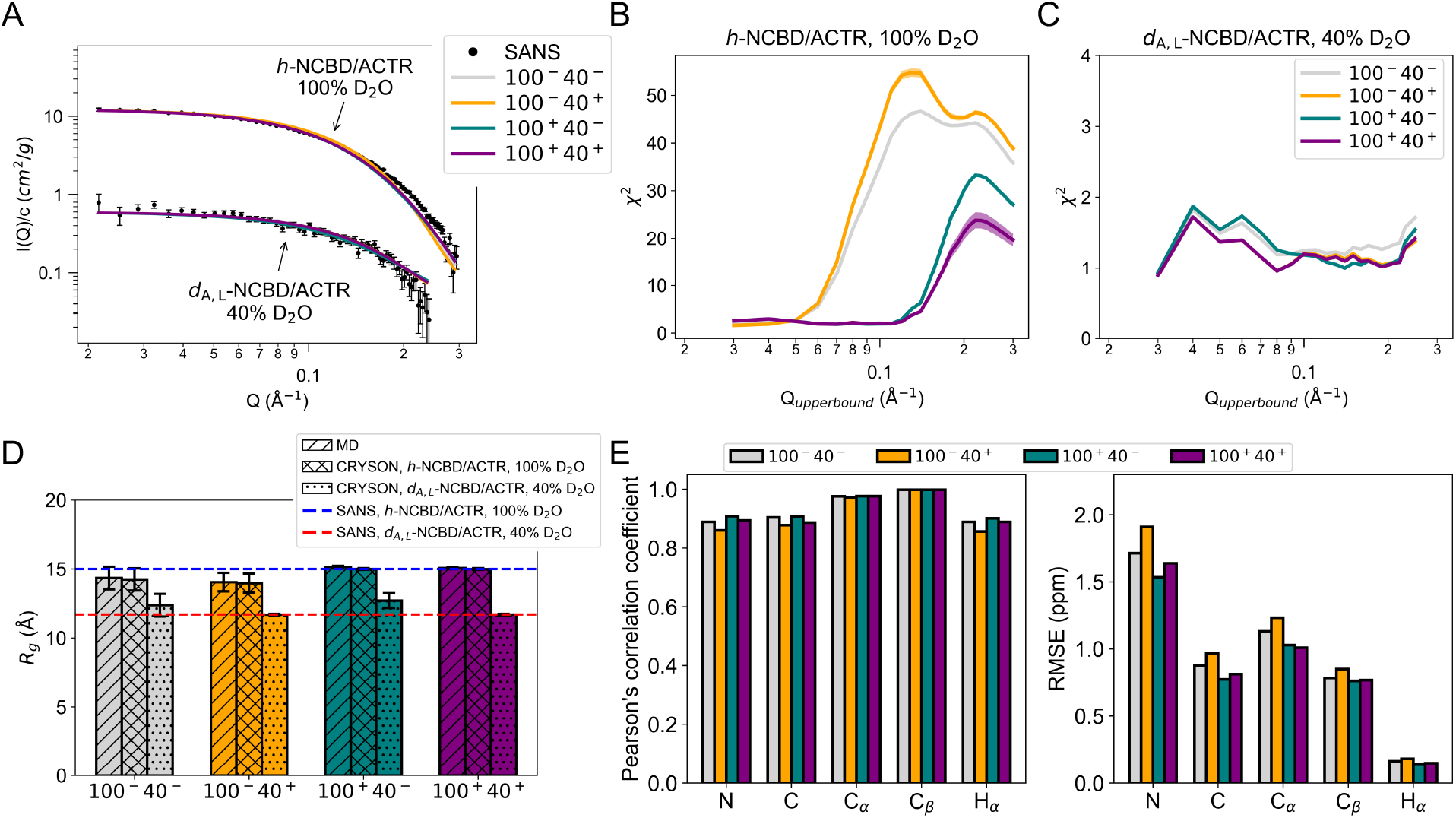
Comparison of NCBD/ACTR *R*_*g*_–based structural ensembles reveals that 100_+_40_+_ has the best agreement to SANS experiment while all *R*_*g*_–based ensembles have better agreement to NMR experiment than the force field–based ensembles. (A-C) Comparison of calculated average SANS curves by CRYSON to experimental SANS curves for 100^-^40^-^, 100^-^40^+^, 100^+^40^-^, 100^+^40^+^ structures. (A) SANS curves normalized by concentration, I(Q)/*c*, against Q from 0.02 Å^−1^ to 0.30 Å^−1^ for *h*-NCBD/ACTR in 100% D_2_O and against Q from 0.02 Å^−1^ to 0.25 Å^−1^ for *d*_A,L_-NCBD/ACTR in 40% D_2_O. The experimental SANS data are shown as black points, where error bars represent the standard deviation. The computed average SANS curves are depicted as lines, where 100^+^40^+^ is in purple, 100^+^40^-^in teal, 100^-^40^+^ in yellow, and 100^-^40^-^in grey. Note that the standard deviations of the computed SANS curves are shown but lie within the line plots. (B, C) Reduced chi-square values (χ^2^) of the SANS curves between experiments and each of the *R*_*g*_–based ensembles for (B) *h*-NCBD/ACTR in 100% D_2_O and (C) *d*_A,L_-NCBD/ACTR in 40% D_2_O. The χ_2_ is plotted against increasing Q range, in which the lower bound is 0.02 Å^−1^ and the upper bound (Q_upperbound_) is from 0.03 Å^−1^ to 0.30 Å^−1^ in B and to 0.25 Å^−1^ in C, with an interval of 0.01 Å^−1^. Shaded error bars represent 95% confidence interval of the mean. Note that the error bars of the χ^2^ for the ensembles are shown but most of them lie within the line plots. (D) Radius of gyration (*R*_*g*_) is computed using atomic coordinates and masses in GROMACS (MD) as well as using CRYSON with and without selective deuteration (CRYSON *d*_A,L_-NCBD/ACTR in 40% D_2_O vs. CRYSON, *h*-NCBD/ACTR in 100% D_2_O). The *R*_*g*_ values derived from the experimental SANS curves of *h*-NCBD/ACTR in 100% D_2_O and *d*_A,L_-NCBD/ACTR in 40% D_2_O are shown as dashed lines in blue and red, respectively. Error bars represent the standard deviation among the structures of each *R*_*g*_–based ensemble. (E) Comparison of computed and experimental chemical shifts by (left) Pearson’s correlation coefficients and (right) RMSEs. Chemical shifts are computed using SPARTA+. These analyses are performed on 233 100^+^40^+^ structures, 6772 100^+^40^-^ structures, 5265 100^-^40^+^ structures, and 95750 100^-^40^-^structures.

Due to varying *R*_*g*_ distribution of MD structures, we seek to gain more structural insights into NCBD and ACTR interactions. To this end, we determined the free energy surface as a function of two structural metrics, the contact area, *A*, between the NCBD and the ACTR and the crossing angle^44, 45^, *θ*, between their longest helices, helix 3 of the NCBD and helix 1 of the ACTR (**Figure 4**). We used all NMR and MD structures to construct a two-dimensional histogram of the normalized probability of *A* and *θ,P/(A, θ)*. Following a similar method used in previous studies^46, 47^, we computed the potential of mean force (PMF) from *W (A, θ)* = −*RT* ln *P (A, θ)*. with a uniform reference distribution. **Figure 4A** highlights the structures satisfying at least one of the experimental *R*_*g*_ constraints from NMR and each force field on the free energy surface. We found the surface has a free energy minimum of −5.2 kcal/mol and is located at *A* = 33 nm^2^ and *θ* = −28°. The negative sign of *θ* denotes that the near helix, i.e., helix 1 of the ACTR, is rotated clockwise relative to the far helix, i.e., helix 3 of the NCBD. The three 100^-^40^+^ NMR structures are in the free energy basin and are overlapped with some structures sampled by the a99SB force field. This similar distribution of *A* and *θ* of the NMR and a99SB structures again suggests the influence of the force field applied in NMR structural refinement. In comparison, the few *R*_*g*_-satisfying a99SB-ILDN structures have a wide distribution, especially in *θ*. Some structures are populated around another local free energy minimum of 0 = 34 nm^2^ and 2 = 38°. As compared with the structures sampled by the a99SB force field, the a99SB-*disp* and C36m structures are populated around the free energy basin with *A* mostly less than 33 nm^2^. The smaller *A* suggests that the NCBD and ACTR of these structures are more flexible and less compact than the structures sampled by the a99SB force fields. We further compared the structures closest to the free energy minimum from each force field. Even with similar values of *A* and *θ*, there are clear differences between these structures (**Figures 4B-4F**). The a99SB and a99SB-ILDN structures have less defined helical content, which is consistent with the helical fraction analysis shown in **Figure S3A**. However, this secondary structural change does not reflect to the same extent on the two reaction coordinates investigated. Compared with the NMR and a99SB-*disp* structures, helix 1 of the ACTR in the C36m structure tilts away as the shape of the accommodating groove formed by helices 1 and 3 of the NCBD shift. In addition, based on the helical content, the ACTR in a99SB-*disp* and C36m structures both gain a helical turn between helices 1 and 2, while the NCBD in a99SB-*disp* structure loses a turn at the end of helix 3. These subtle global and local structure details are indistinguishable on the free energy surface but become apparent when visualizing their three-dimensional (3D) structures. From the distribution of different structural ensembles on the free energy surface, we observe distinct regions sampled by different force fields. Nevertheless, using only these two structural metrics is insufficient to distinguish finer structural details.

**Figure 4.**
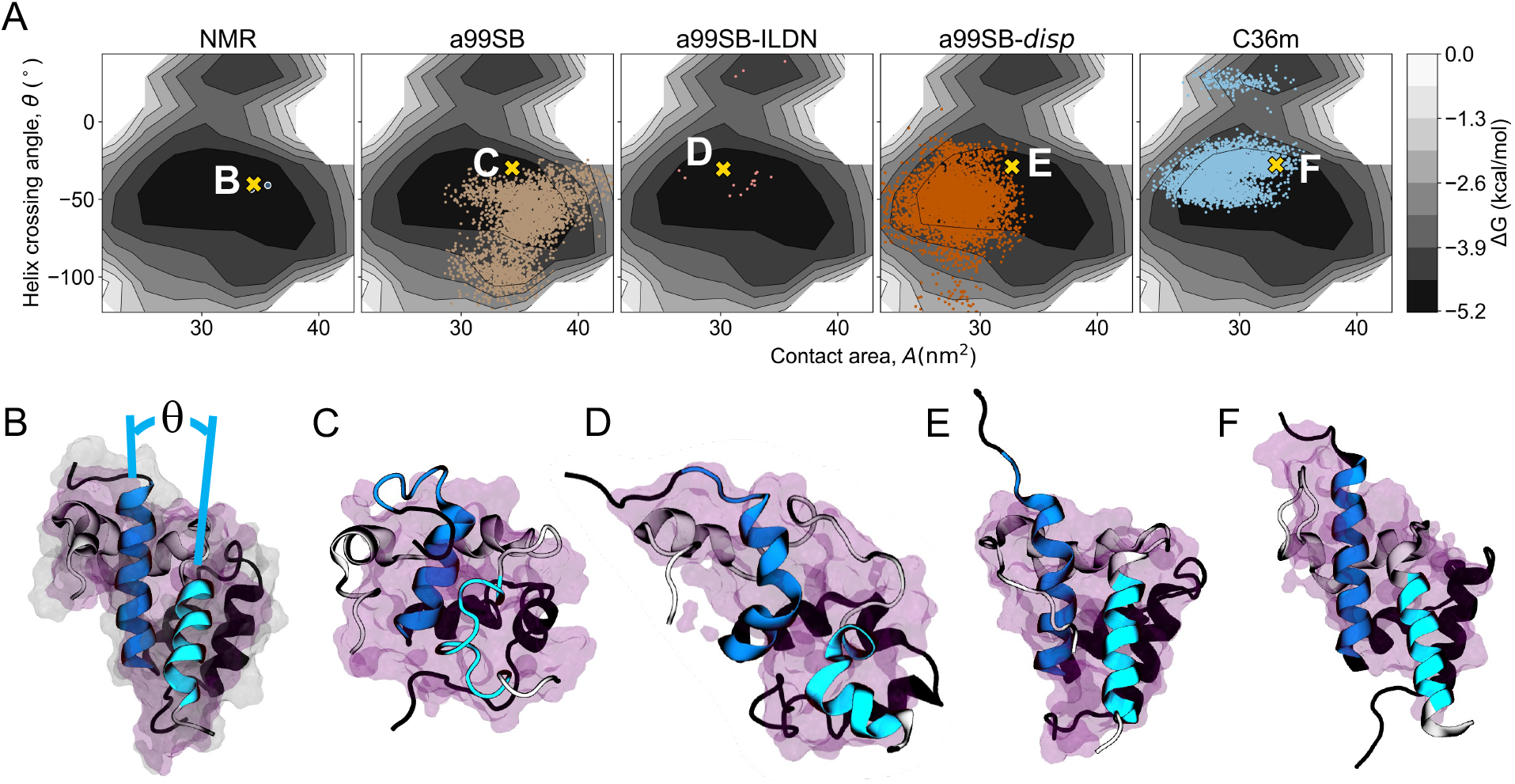
Representative NMR and MD structures are structurally different yet indistinguishable on the free energy surface. (A) Free energy surface of all NMR and MD structures in terms of two reaction coordinates, the contact area, *A*, between NCBD and ACTR and the crossing angle, *θ*, between their longest helices. Each panel shows the NMR and MD structures from each force field satisfying at least one *R*_*g*_ constraints from SANS experiments. Data points are color coded as in Figure 2. An NMR structure and a representative MD structure closest to the free energy minimum from each force field is highlighted by a yellow ‘x’ and presented in B, C, D E, and F, respectively. (B-F) Representative NCBD (black)/ACTR (gray) structures selected from the free energy surface. The longest helices of NCBD and ACTR are colored in blue and cyan, respectively. The contact area is depicted by magenta surface. The rest of the complex surface is shown in light grey in the NMR structure but omitted in the other structures for clarity.

To comprehensively compare the NCBD/ACTR complex structures, we investigated an alternative method using a deep learning (DL) technique. We selected the NMR and MD structures satisfying at least one of the experimental *R*_*g*_ constraints (i.e., all data points shown on the free energy surface in **Figure 4A**) and applied a convolutional variational autoencoder (CVAE) to encode high dimensional protein complex structures into a 3-D latent space for visualization. The CVAE is a variational autoencoder (VAE)^48^ where the encoder and decoder are convolutional neural networks. The encoder of the CVAE serves as a dimensionality reduction tool which effectively projects high dimensional molecular structures in a lower dimensional, normally distributed latent space, in which similar structures are placed close to one another^49, 50^. The decoder, which is symmetric but in reverse order to the encoder, reconstructs the input from sampling of the constructed latent space. Direct comparison of the decoded data with the original input ensures accuracy of the latent space representation. Together, the loss function of the CVAE is a sum of the Kullback-Leibler divergence of the encoder output distribution from the standard normal distribution and the reconstruction loss between the decoder output and the original input. By minimizing the loss function, the CVAE model learns to compress and reconstruct data between high dimensional input space and a low dimensional representation while maintaining high integrity^51^. This feature allows efficient data analysis and data visualization by comparing different complex molecular structures in the low dimensional latent space. Decision on the latent space dimension (i.e., 3-D) was mainly for visualization purpose and for selecting the lowest possible dimension that can effectively represent the input without sacrificing the loss. The CVAE has been successfully applied to study the folding pathways of small proteins^52^ and structural clustering of biomolecules^53-55^.

We represented each *R*_*g*_-satisfying structure by a distance matrix calculated between the C_*α*_ atoms of every residue pair. Distance matrix representation removes translational and rotational variances in the 3-D molecular structures. The CVAE model learned structural features presented in the distance matrices and projected similar structures close to each other in the latent space. The latent space reveals separation between MD structures sampled by different force fields (**Figure 5A**), suggesting distinct structural features between different MD ensembles. Particularly, the structures sampled by the a99SB and a99SB-*disp* force fields are projected away from those by the a99SB-ILDN and C36m force fields, which agrees with the overall MD structure distribution on the free energy surface.

**Figure 5.**
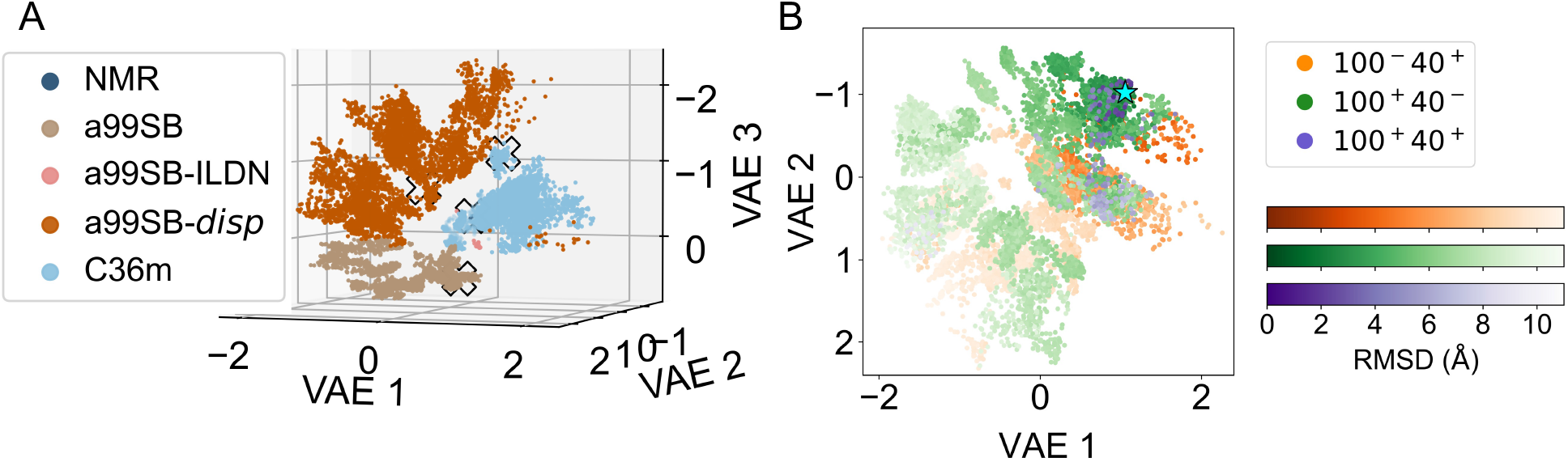
A convolutional variational autoencoder model discerns structural differences in the NCBD/ACTR structures from NMR spectroscopy and MD simulations. (A) The 3-D latent space representing a total of 9,816 NMR and MD structures in the training set. Clusters are labeled by the experimental method and force fields and color coded as in Figure 2. Note that some C36m data are dimmed to reveal the NMR data and the y- and z-axes are reversed due to the viewing angle of the 3-D plot. White ‘x’ marks are the representative NMR, a99SB, a99SB-*disp*, and C36m structures selected from the free energy surface in Figure 4, showing clear separation between these structures in the CVAE latent space. The representative a99SB-ILDN structure is in the validation set and therefore not on the graph. (B) The 3-D latent space projected onto the VAE 1-VAE 2 plane. Structures are labeled based on conditional agreement of *R*_*g*_ values from SANS experiments and all-atom RMSD values with respect to a reference 100^+^40^+^ structure highlighted by a cyan star. Structures that satisfy either one of the *R*_*g*_ constraints (100_-_40_+_ or 100_+_40_-_) are in yellow and teal, respectively. Structures that are within both constraints (100^+^40^+^) are in purple.

To elucidate structural features that the model learned to encode and decode, we labeled the same latent space by the *R*_*g*_ categories defined previously and projected the space in a 2-D plane with the structures labeled by their all-atom RMSD values with respect to a 100^+^40^+^ structure as the reference (**Figure 5B**). Interestingly, the 100^+^40^+^ structures are in the regions in which the 100^-^40^+^ and 100^+^40^-^structures overlap, and there is a clear trend that the RMSD value increases as the structure is projected further away from the reference. These results further demonstrate that the CVAE model learns the structural details embedded in the distance matrices and the resulting latent space presents a road map that allows comprehensive comparison of different NCBD/ACTR complex structures. Compared to commonly applied structural analysis methods, which are mostly limited to specific regions of the structure of interest, this DL approach allows us to evaluate the structure as a whole and still capture detailed local structural features^55^. Moreover, unlike other dimensionality reduction methods which rely on pre-defined collective variables, the CVAE constructs the underlying low dimensional representation space in an unsupervised manner.

To further characterize the 100^+^40^+^ structures, we analyzed the projection of their distance matrices in the latent space. The structures arrange approximately into three clusters described by RMSD ranges, (1) from 0 to 5 Å, (2) from 5 to 8 Å, and (3) from 8 Å and above (**Figure 6A**), using a 100_+_40_+_ structure as the reference. The C_*α*_ RMSF values of the ACTR and NCBD structures demonstrate similar structural fluctuations between the three clusters. In addition to the unstructured regions at the N- and C-terminal ends of both proteins, which fluctuate up to 12.7±2.5 Å, there are three internal regions of the complex varying between 2.8 Å and 4.2 Å (**Figure 6B**). These flexible regions roughly correspond to lower average helical fraction of the complex. However, there are various helical contents, especially in ACTR, with low fluctuation between the three clusters (**Figure 6D**). The contact area between NCBD and ACTR of the 100^+^40^+^ structures is less than 30 nm^2^ and the helix crossing angle around −50°. **Figure 6C** shows their location on the free energy surface. When highlighting the flexible regions on a 3-D NCBD/ACTR complex structure, we found that they are located at the junctions connecting neighboring helices (**Figure 6E**). One flexible region consisting of ACTR residues 1057 to 1063 connects helices 1 and 2. The other two flexible regions are in the NCBD. Residues 2077 to 2085, which form a part of the polyglutamine tract, connect helices 1 and 2, while residues 2092 to 2097 connect helices 2 and 3. These flexible regions are in agreement with observations from a previous NMR relaxation study^30^. Representative complex structures selected from the three clusters reveal distinct orientation and organization of the helices (**Figures 6F-6H)**. Based on the number of structures in each cluster, we calculated their composition percentage in the 100^+^40^+^ ensemble, which are 37 %, 49 %, and 14 %, respectively. Globally, the three helices of the ACTR display different tilting angles with respect to the NCBD. Locally, structural flexibility in the NCBD reveals that a helical turn in the polyglutamine tract and regions near the two terminal ends may become random coils. In summary, our SANS-derived *R*_*g*_ and PMF analysis suggests that the structural ensemble of the NCBD/ACTR complex is heterogenous, in which distinct complex arrangements are in good agreement with SANS experiments. Nevertheless, from our integrated method of SANS and MD/DL, we have characterized a structural ensemble which describes SANS and NMR experiments better than other ensembles determined by MD simulation with one force field. Although obtaining sufficient neutron scattering signal is a main driver in designing residue-based selective labeling of SANS, the flexibility of labeling choice in combination with varying contrast of D_2_O concentration in the solvent offers an auspicious option to study structurally flexible systems such as IDPs. Selective labeling and contrast matching SANS experiments provide not only shape and size as constraints to select from an initial pool of conformations but also average scattering intensity for more stringent validation.

**Figure 6.**
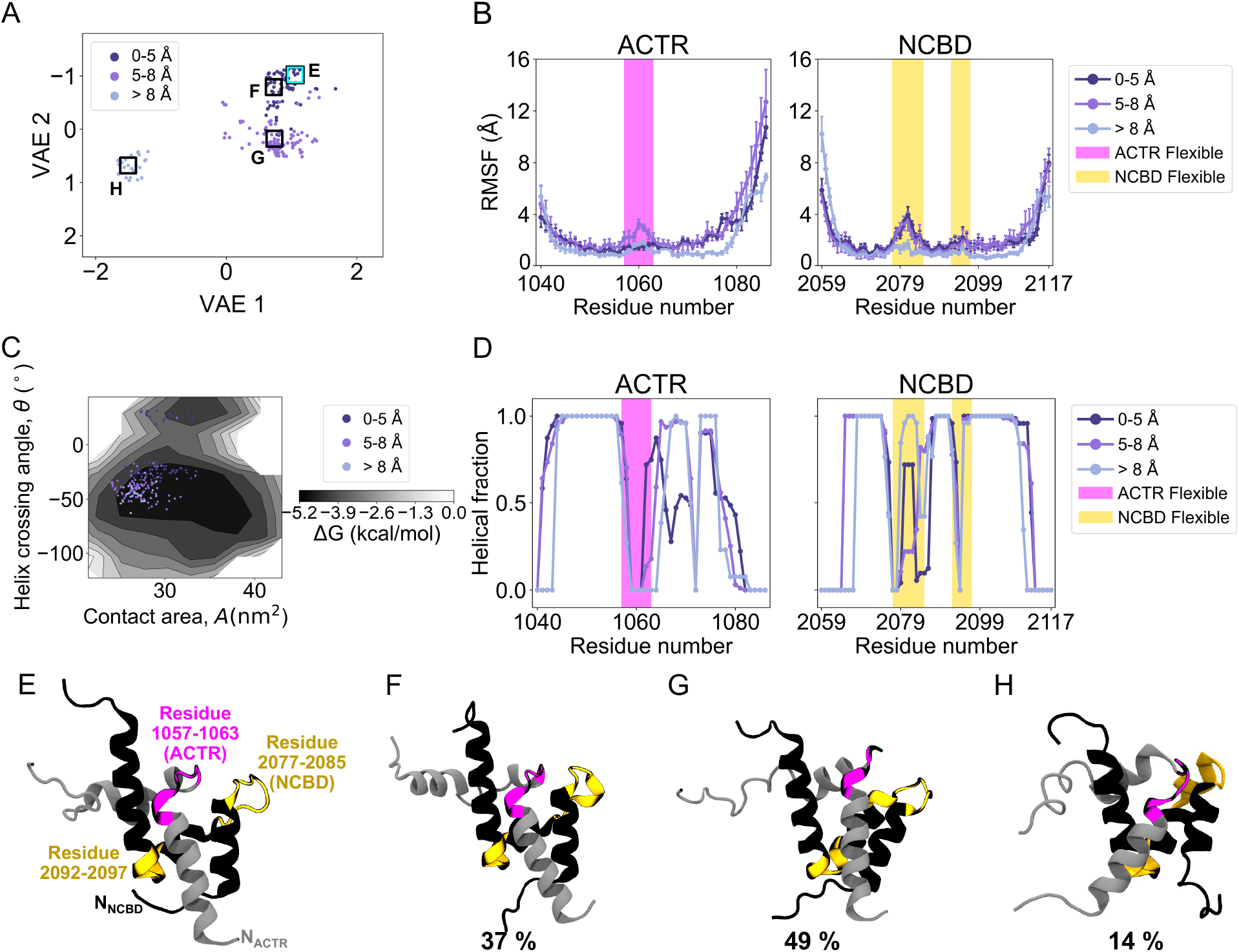
Characterization of NCBD/ACTR structural ensemble based on the structures satisfying both experimental *R*_*g*_ constraints reveals three distinct clusters. (A) The 3-D latent space projected onto the VAE 1-VAE 2 plane with only the 100^+^40^+^ structures. Structures are labeled based on all-atom RMSD values using cutoff distances of 5 Å and 8 Å with respect to the same reference 100^+^40^+^ structure shown in Figure 5B highlighted by a cyan box and presented in E. The structure closest to the centroid of each cluster is boxed and presented in F, G, and H, respectively. (B) The C_*α*_ RMSF values of ACTR and NCBD structures in the three clusters. The flexible region of ACTR is highlighted in magenta while that of NCBD is in yellow. Error bars represent the standard deviation. (C) The 100^+^40^+^ structures on the free energy surface shown in Figure 4A. (D) Helical fraction of 100^+^40^+^ structures. The flexible regions are highlighted again for comparison. (E) Reference 100^+^40^+^ structure for RMSD. (F-H) Representative NCBD (black)/ACTR (gray) structures selected from the three clusters, each with its cluster’s composition percentage in the ensemble. The flexible regions are color coded as in B and D.

## Conclusion

In this work, we characterize the structural ensemble of an intrinsically disordered protein complex consisting of the NCBD and ACTR domains by small-angle neutron scattering (SANS) and computing. Beyond full contrast SANS experiments, residue-specific deuterium labeling with contrast matching presents additional information of the same system to aid characterization of structural ensembles. We combine SANS experiments with molecular dynamics (MD) simulations using four different AMBER and CHARMM force fields and a deep learning (DL) convolutional variational autoencoder (CVAE) to explore the structural space of the NCBD/ACTR complex. We find that each force field generates a distinct pool of structures, where the a99SB-*disp* and C36m structural ensembles show better agreement with the experimental SANS and NMR data. Applying structural constraints, such as the radius of gyration (*R*_g_) from SANS experiments, to classify structures leads to a structural ensemble that fits the SANS scattering curves and the NMR chemical shifts better than other ensembles generated by MD simulations with one force field. Complementary to structural metrics like contact area or helix crossing angle, the CVAE algorithm allows for comprehensive comparison of complete three-dimensional structures using their distance matrices by projecting similar structures closer in a lower-dimensional latent space. From structure projection in the latent space, we characterize an *R*_g_-based ensemble, determining three representative conformations and the corresponding composition percentage. Taken together, we present an integrated SANS and MD/DL method for characterizing the structural ensemble of an intrinsically disordered protein complex. This work provides insights for further study of more structurally flexible systems.

## Materials and Methods

### I. Sample preparation

The nuclear receptor coactivator binding domain (NCBD) of mouse cAMP response element binding (CREB) protein (CBP, accession: NP_001020603), CBP(2059-2117), and the interaction domain of mouse activator for thyroid hormone and retinoid receptor (ACTR, accession: Q9Y6Q9), ACTR(1046-1093) were synthesized by solid-phase FMOC chemistry and purified to >95%. Hydrogenated NCBD (*h-*NCBD, 6545 Da) was synthesized by Keck Yale facility. Selectively deuterated NCBD (*d*_A,L_-NCBD, 6633 Da) was synthesized by New England Peptide with L-alanine-N-Fmoc (2,3,3,3-D4, 98%) and L-leucine-N-Fmoc (D10, 98%) to label all five alanine and seven leucine residues. Hydrogenated ACTR (5214 Da) also was synthesized by New England Peptide. Mass spectrometry confirmed the molecular mass of the peptides. D_2_O (99.9% D) (Cambridge Isotope Laboratories, Inc. Tewksbury, MA), monosodium phosphate, disodium phosphate (Sigma-Aldrich), and sodium chloride (Fluka) were used without further purification.

### II. Small-angle neutron scattering

SANS experiments were performed on the extended Q-range small-angle neutron scattering beam line at the Spallation Neutron Source located at Oak Ridge National Laboratory^31^. In 60 Hz operation mode, a 2.5 m sample-to-detector distance with 2.5-6.4 Å wavelength band was used to obtain the relevant wavevector transfer, *Q* = 4*π* sin(*θ*)/*A*, where 2*θ* is the scattering angle. NCBD/ACTR samples were prepared in 20 mM sodium phosphate (pH 7), 50 mM NaCl, H_2_O/D_2_O. *h*-NCBD/ACTR (2.5 mg/mL) in 100% D_2_O, *h*-NCBD/ACTR (10.3 mg/mL) in 40% D_2_O, and *d*_A,L_-NCBD/ACTR (8.7 mg/mL) in 40% D_2_O buffer were measured. Peptide concentrations were determined by UV-Vis using a calculated absorption, 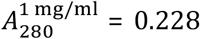 for NCBD^56^. Samples were loaded into either 1 or 2 mm pathlength circular-shaped quartz cuvettes (Hellma USA, Plainville, NY) and SANS measurements were performed at 20 °C. Data reduction followed standard procedures using MantidPlot^57^. The measured scattering intensity was corrected for the detector sensitivity and scattering contribution from the solvent and empty cells, and then placed on absolute scale using a calibrated standard^58^.

### III. SANS analysis

Guinier fits to the low-*Q* region of the *I*(*Q*) scattering curves were initially performed to confirm a Guinier regime^59^ (**Figure S1**). The pair distance distribution function, *P*(*r*), was then calculated from the *I*(*Q*) curves using the GNOM program^60^ (**Figure 1C**). The *P*(*r*) function was set to zero for *r* = 0 and *r* = *D*_max_, the maximum linear dimension of the scattering object, and the *P*(*r*) was normalized to the peak maximum in each profile. The real-space radius of gyration, *R*_g_, and scattering intensity at zero angle, *I*(0), were determined from the *P*(*r*) solution to the scattering data. Molecular mass, *M*, was calculated from

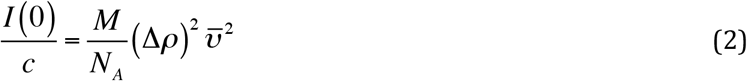

where Δ*ρ* = contrast in scattering length density between protein and D_2_O buffer solution (= *ρ*_prot_ – *ρ*_buf_), 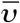 partial specific volume and *N*_A_ = Avogadro’s number. The protein scattering length density, *ρ*_prot_, of NCBD/ACTR complexes were calculated from the sequence using the Contrast module of MULCh_32_. The D_2_O scattering length density used was *ρ*_D2O_ = 6.388 × 10_10_ cm_-2_. MULCh calculations yielded Δ*ρ* = –3.298, 0.098, and 0.75 × 10_10_ cm_-2_ for *h*-NCBD/ACTR in 100% D_2_O, *h*-NCBD/ACTR in 40% D_2_O, and *d*_A,L_-NCBD/ACTR in 40% D_2_O, respectively. The MULCh value for 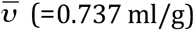 of NCBD/ACTR also was used in the above equation. Experimental SANS values are given in **Table S1**.

### IV. Molecular systems and MD simulations

We performed all-atom MD simulations to determine the dynamic structure of the complex formed by the two intrinsically disordered proteins, NCBD and ACTR. We investigated four state-of-the-art force fields from Amber and CHARMM families, including a99SB^35^, a99SB-ILDN^36^ with TIP3P water model^61^, a99SB-*disp*^37^, and C36m with the CHARMM-modified TIP3P water model^38^. Both a99SB-*disp* and C36m force fields were recently developed to address structural flexibility and improve accuracy of disordered proteins in simulations. We used an NMR structure of the NCBD/ACTR complex in PDB 1KBH^29^ as the initial structure. The complex consists of a total of 106 residues, in which the first 47 residues are in chain A, which belongs to ACTR, and the remaining 59 residues are in chain B, which is part of NCBD. The complex structure was solvated in the center of a water box with a minimum distance of 15 Å from the edge of the box to the nearest protein atom, neutralized with counter ions and ionized with 50 mM NaCl following the experimental conditions. We then minimized and equilibrated the resulting system for 100 ns, followed by a 1 µs trajectory in a production run. For each force field, we performed three independent 1 µs trajectories. After the first 100 ns, we sampled structures every 100 ps for analysis, yielding a total of 27,000 structures for each force field.

All MD simulations were performed with OpenMM^62^ in NPT ensemble at 1 atm and 293.15 K with a time step of 2 fs. Nonbonded interactions were calculated with a typical cutoff distance of 12 Å, while the long-range electrostatic interactions were enumerated with the Particle Mesh Ewald algorithm^63^.

### V. Structure assessment by CRYSON^41^

To evaluate the 108,000 NCBD/ACTR complex structures from MD simulations and the 20 NMR models in PDB 1KBH^29^, we used the CRYSON program^41^ to compute SANS curves using their atomic structures and compared these curves with scattering data from SANS experiments. Following the experimental conditions, for each structure we calculated two scattering curves, one with hydrogenated NCBD/ACTR complex (*h*-NCBD/ACTR) in 100% D_2_O, and the other with selectively deuterated NCBD/ACTR complex (*d*_A,L_-NCBD/ACTR) in 40% D_2_O. For *d*_A,L_-NCBD/ACTR in 40% D_2_O, all five alanine and seven leucine residues were labeled as a separate chain (i.e., chain C) in their atomic coordinate files and set to fraction deuterated of 1 in CRYSON. All other residues in the original chains (i.e., chain A and chain B) were set to fraction deuterated of 0. The fraction of D_2_O in the solvent was set to 0.4.

All CRYSON calculations were performed with the maximum scattering vector of 0.3 Å^−1^ and 200 points in the scattering curve. Explicit hydrogen atoms were considered, and the contrast of the solvation shell was set to 0. No fitting to experimental data was involved in the scattering curve calculations to provide unbiased comparison between the theoretical and experimental scattering curves. The experimental *R*_g_ values serve as the constraints to select qualified MD and NMR structures. For each structure, we derived *R*_g_ values from the two scattering curves. Only the structures satisfying at least one of the *R*_g_ constraints from SANS experiments were included for deep learning analysis. The total number of *R*_g_-satisfying structures was 12,270, which was about 11.4% of 108,020 structures generated from MD simulations and NMR spectroscopy.

### VI. Deep learning analysis

To systematically analyze the *R*_g_-satisfying NCBD/ACTR complex structures, we deployed an autoencoder-based deep learning architecture, a convolutional variational autoencoder (CVAE), for structure comparison and visualization at large scale. To generate translation and rotation invariant input data for CVAE, we represented each *R*_g_-satisfying structure by a distance matrix using the C_*α*_ atoms of the protein complex. There are 106 residues in the NCBD/ACTR complex, so the size of each distance matrix was 106 × 106. We then merged the distance matrices of the 12,270 MD and NMR structures and randomly split the matrices into training and validation datasets using the 80/20 ratio. We then constructed a CVAE model to project these high dimensional structural data into a 3-D latent space using the *Keras* Library^64^. The encoder model consisted of three convolutional layers and a fully connected layer, each with 64 feature maps. We used a 2 × 2 convolution kernel and a stride of 1, 2, and 1 at the three convolutional layers, respectively. The activation function at each convolutional layer was ReLu. The optimizer was RMSProp^65^, with a learning rate of 0.001. We trained the model for 250 epochs, along which the training and validation loss converged (**Figure S5**). The difference between decoded and original images is minimal, suggesting the model was trained successfully (**Figure S6**). We then analyzed the structures from the clusters in the latent space and selected representative structures for visualization using VMD^66^.

## Acknowledgements

We would like to thank David Bell and Max Chen for helpful discussions. We would also like to thank Chris Layton and Daniel Dewey for technical support.

This work was performed at the Compute and Data Environment for Science (CADES) of the Oak Ridge National Laboratory (ORNL), which is funded by the Office of Science of the U.S. Department of Energy under Contract No. DE-AC05-00OR22725. A portion of this research was performed at Oak Ridge National Laboratory’s Spallation Neutron Source, sponsored by the U.S. Department of Energy, Office of Basic Energy Sciences. We acknowledge laboratory support by the Center for Structural Molecular Biology, funded by the Office of Biological and Environmental Research of the U.S. Department of Energy. This research was supported by ORNL Laboratory Directed Research and Development SEED grant No. 7278.

The research was supported by the U.S. Department of Energy, Office of Science, Office of Advanced Scientific Computing Research, under contract number DE-AC05-00OR22725; the Joint Design of Advanced Computing Solutions for Cancer (JDACS4C) program established by the U.S. Department of Energy (DOE) and the National Cancer Institute (NCI) of the National Institutes of Health. It was performed under the auspices of the U.S. Department of Energy by Argonne National Laboratory under Contract DE-AC02-06-CH11357, Lawrence Livermore National Laboratory under Contract DE-AC52-07NA27344, Los Alamos National Laboratory under Contract DE-AC5206NA25396, Oak Ridge National Laboratory under Contract DE-AC05-00OR22725, and Frederick National Laboratory for Cancer Research under Contract HHSN261200800001E.

## Author contributions

S.H.C. designed the study and performed MD simulations and data analysis. K.L.W. and C.S. designed and performed SANS experiments. S.H.C. and D.B. developed the CVAE code. S.H.C. and C.S. wrote the paper. All authors read and approved the final manuscript.

## Materials and correspondence

Correspondence and requests for materials should be addressed to S.H.C.

## Availability of materials and data

All relevant data are available from within the manuscript and Supplementary Information, and from the corresponding author upon reasonable request.

## SUPPORTING INFORMATION

**Figure S1.**
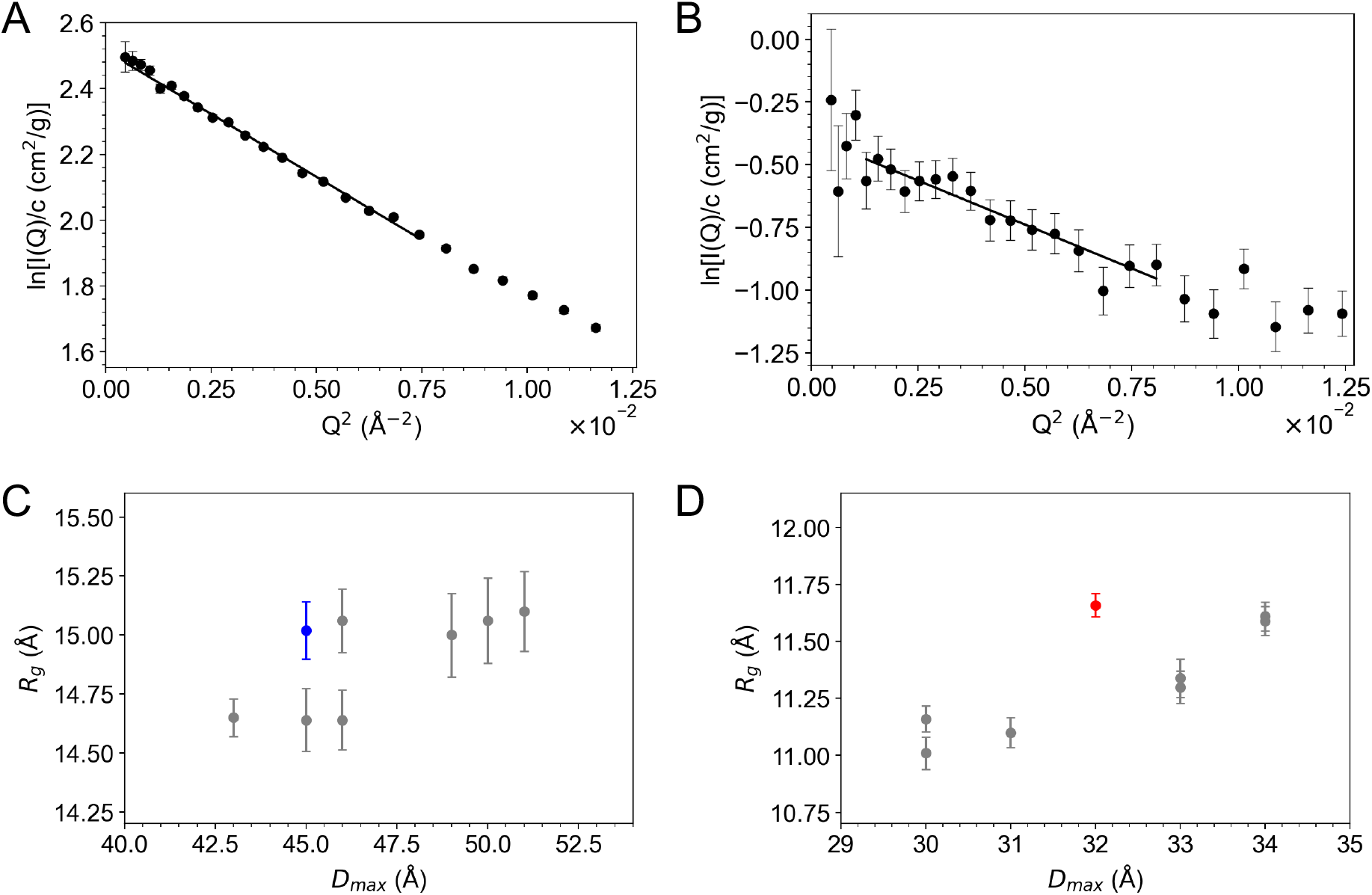
Guinier fits and *R*_g_–*D*_*max*_ correlation reveal consistent *R*_g_ values. Guinier fits performed on concentration-normalized scattering data, *I*(*Q*)/*c*, for (A) *h*-NCBD/ACTR in 100% D_2_O and (B) *d*_A,L_-NCBD/ACTR in 40% D_2_O. The calculated *I*(0)/*c* and *R*_g_ values are given in **Table S1**. The *R*_*g*_ vs. *D*_*max*_ values obtained from multiple *P*(*r*) fits to the scattering data for (C) *h*-NCBD/ACTR in 100% D_2_O and (D) *d*_A,L_-NCBD/ACTR in 40% D_2_O (as shown in Figure 1C). The values for each that are used for all further analyses are highlighted (blue and red symbols, respectively).

**Table S1.** NCBD/ACTR experimental SANS values.

| | Guinier | | $P(r)$ | | | |
| --- | --- | --- | --- | --- | --- | --- |
| | $I(0)/c$ ($cm^2/g$) | $R_g$ ($\text{\AA}$) | $I(0)/c$ ($cm^2/g$) | $M$ (kDa)<sup>1,2</sup> | $R_g$ ($\text{\AA}$) | $D_{max}$ ($\text{\AA}$) |
| $h$-NCBD/ACTR, 100% $D_2O$ | $12.36 \pm 0.07$ | $15.2 \pm 0.1$ | $12.2 \pm 0.1$ | $12.4 \pm 0.1$ | $15.0 \pm 0.1$ | 45 |
| $d_{A,L}$-NCBD/ACTR, 40% $D_2O$ | $0.68 \pm 0.02$ | $14.5 \pm 0.7$ | $0.596 \pm 0.004$ | $11.74 \pm 0.09$ | $11.7 \pm 0.1$ | 32 |

<sup>1</sup> $M$ calculated from $P(r)$ determined $I(0)/c$ values using eq. (2) in main text.
<sup>2</sup>Actual $M = 11.9$ kDa for NCBD/ACTR.

**Figure S2.**
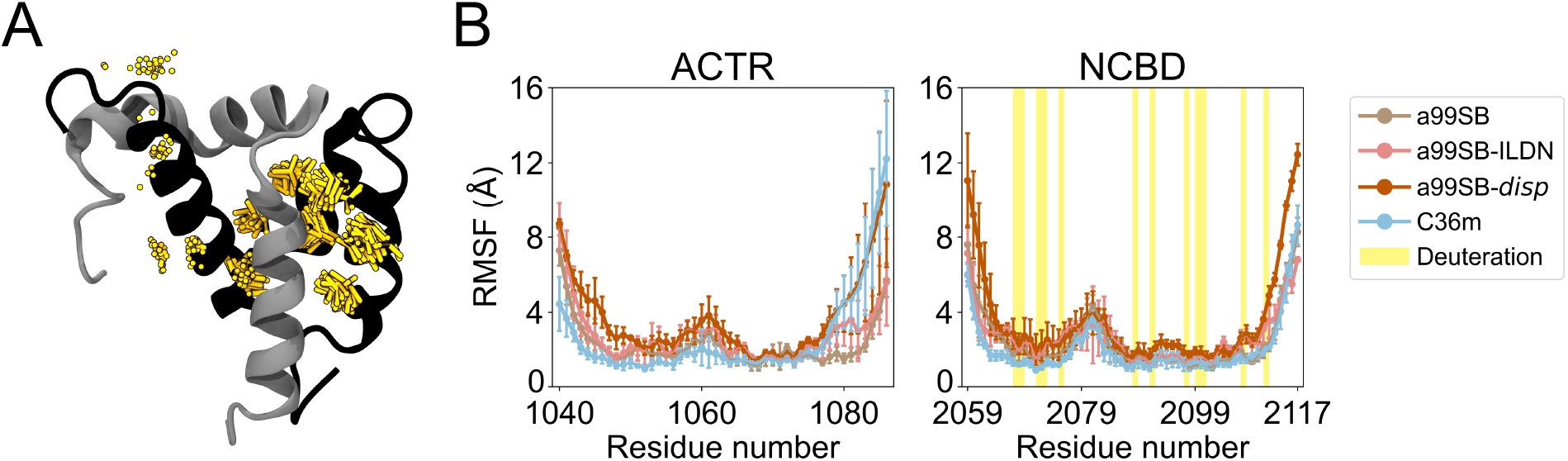
Deuterated positions are relatively stable along MD trajectories. (A) A representative trajectory with the C36m force field. NCBD and ACTR structures are in black and gray ribbons, respectively. NCBD was selectively deuterated at all five Ala and seven Leu positions, where the heavy atoms of their side chains are overlayed in yellow sticks along 1 µs, with the first 100 ns discarded. (B) The average C_*α*_ RMSF values of ACTR and NCBD structures in four MD ensembles. For each ensemble, C_*α*_ RMSF values are averaged across three independent trajectories of 9,000 structures each (the first 100 ns/1,000 frames were discarded), for a total of 27,000 structures per ensemble. The deuterated residues are highlighted in yellow. Error bars represent the standard deviation among three independent trajectories of each ensemble.

**Figure S3.**
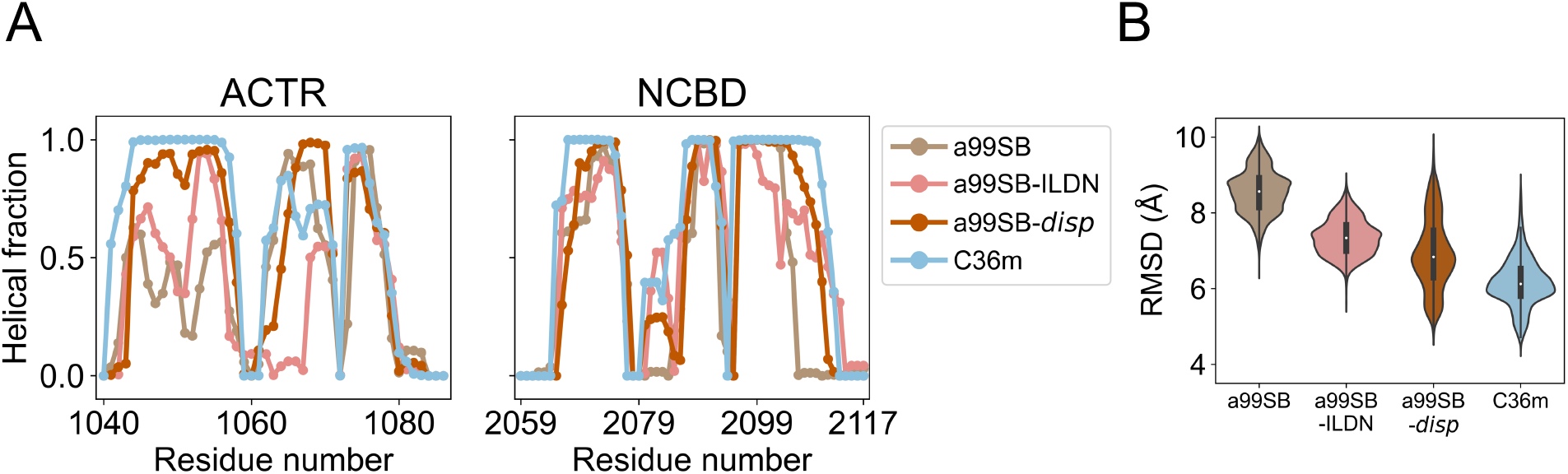
MD structures sampled by different force fields show distinct distributions of helical fraction and root mean square deviation. (A) Helical fraction and (B) all-atom root mean-square deviation (RMSD) of structures sampled from different MD force fields. For each MD ensemble, the values are averaged across 27,000 MD structures taken from three independent trajectories, with the first 100 ns discarded. The reference structure of the RMSD is a solution NMR structure (PDB ID 1KBH). The whiskers represent the 95% confidence intervals.

**Figure S4.**
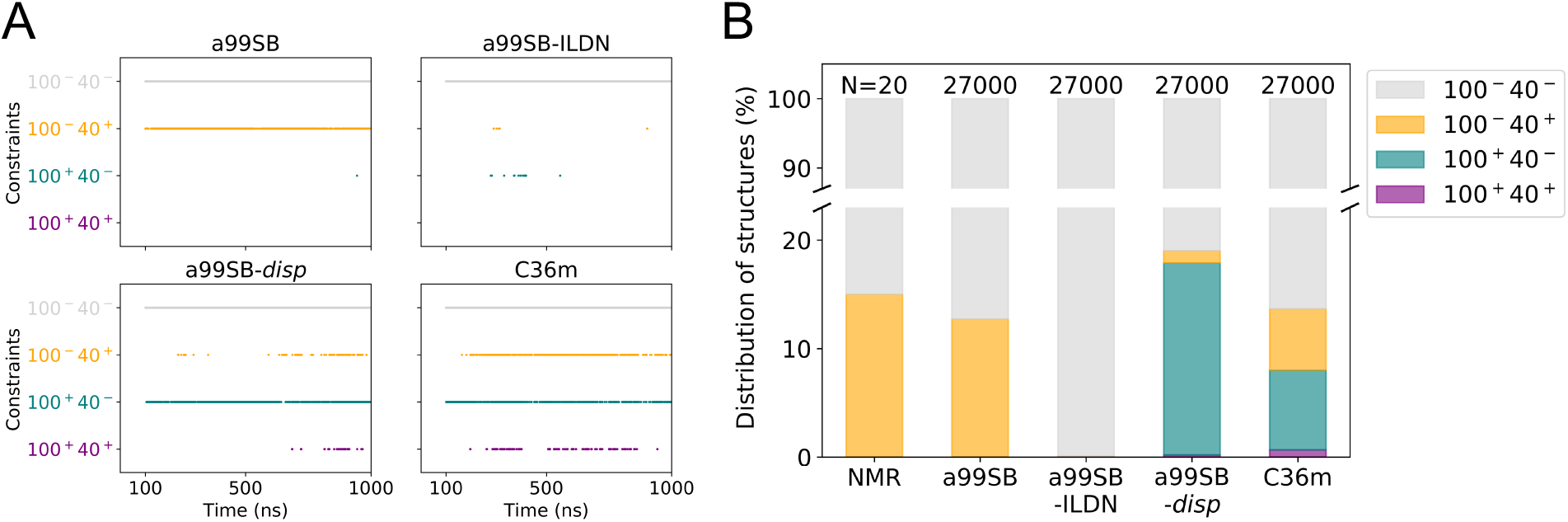
Classification of NCBD/ACTR MD structures based on their *R*_*g*_ values derived by CRYSON. (A) For each of the four force fields, a99SB, a99SB-ILDN, a99SB*-disp*, and C36m, 27,000 MD structures are taken from three independent trajectories, with the first 100 ns discarded. At every given time point, three structures, one from each replica, are plotted. For each structure, the *R*_*g*_ values are calculated under two conditions, *h*-NCBD/ACTR in 100% D_2_O (100) and *d*_A,L_-NCBD/ACTR in 40% D_2_O (40). Structures that do not satisfy either *R*_*g*_ constraints from SANS experiments (100_-_40_-_) are in grey. Structures that satisfy either one of the constraints (100^-^40^+^ or 100^+^40^-^) are in yellow and teal, respectively. Structures that are within both constraints (100^+^40^+^) are in purple. (B) Distribution of the 100^-^40^-^, 100^-^40^+^, 100^+^40^-^, 100^+^40^+^ structures in the NMR and MD ensembles.

**Figure S5.**
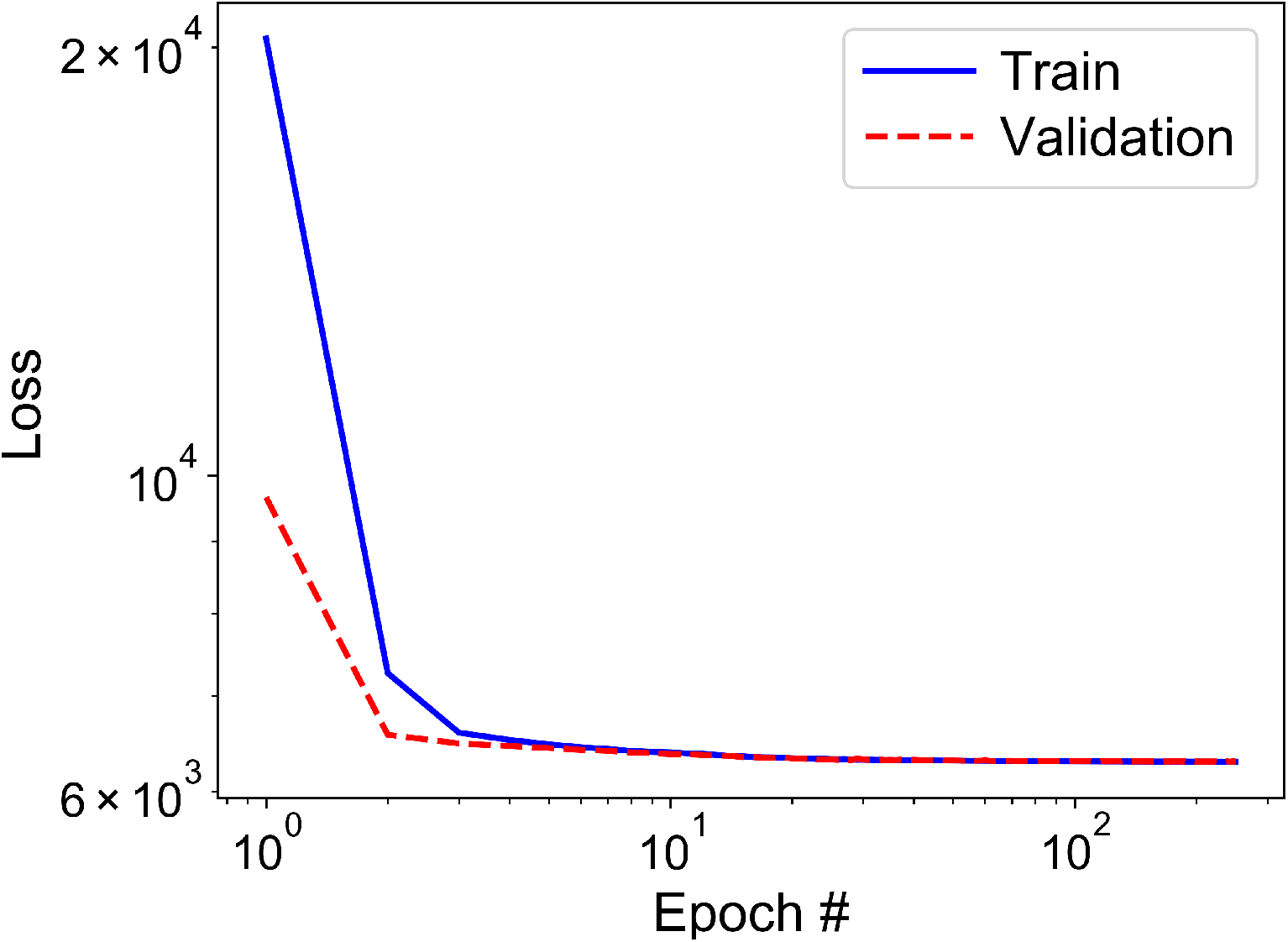
Training and validation losses converge by 250 epochs. The loss curves when training the CVAE model on the distance matrices of NMR and MD structures along 250 epochs. Only the structures satisfying at least one of the *R*_*g*_ constraints from SANS experiments are included for training and validation. Note that both axes are in log scale.

**Figure S6.**
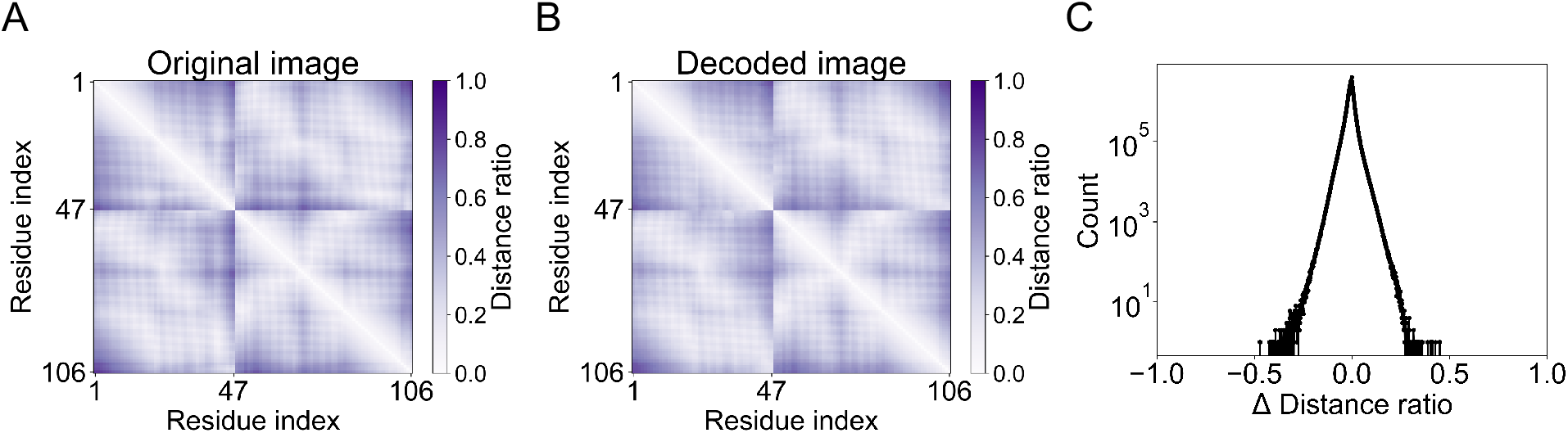
CVAE model is trained to reconstruct distance matrices. Assessment of the CVAE model trained on a total of 9,816 NMR and MD structures after 250 epochs. (A) An example original image, the distance matrix of the reference 100^+^40^+^ structure shown in Figure 6E. Note that the distance range is normalized to 0 and 1. (B) The corresponding decoded image using the trained model. Residue indices 1 to 47 belong to ACTR and 48 to 106 belong to NCBD. (C) Enumerated difference between the decoded and original images in the training set. Note that the y-axis is in log scale.

